# Transcriptional Signatures Associated with the Survival of *Escherichia coli* Biofilm During Treatment with Plasma-Activated Water

**DOI:** 10.1101/2024.05.07.593042

**Authors:** Heema Kumari Nilesh Vyas, M. Mozammel Hoque, Binbin Xia, David Alam, Patrick J. Cullen, Scott A. Rice, Anne Mai-Prochnow

## Abstract

Biofilm formation on surfaces, tools and equipment can damage their quality and lead to high repair or replacement costs. Plasma-activated water (PAW), a new technology, has shown promise in killing biofilm and non-biofilm bacteria due to its mix of reactive oxygen and nitrogen species (RONS), and in particular superoxide. However, the specific genetic mechanisms behind PAW’s effectiveness, especially against biofilms, are not yet fully understood. Here, we examined the stress responses of *Escherichia coli* biofilms when exposed to sub-lethal PAW treatment with and without superoxide (by adding the scavenger Tiron to remove it). A 40% variation in gene expression was observed for PAW treated biofilms when compared to PAW-Tiron and controls. Specifically, PAW treatment resulted in 478 upregulated genes (> 1.5 log2FC) and 186 downregulated genes (< −1.5 log2FC) compared to the control. Pathway enrichment and biological process enrichment analysis revealed significant upregulation of sulfur metabolism, ATP-binding cassette transporter genes, amino acid metabolic/biosynthesis pathways, hypochlorite response systems and oxidative phosphorylation for biofilms treated with PAW compared to control. Knockout mutants of significantly upregulated genes associated with these pathways *trxC* (4.23-fold), *cysP* (1.58-fold) and *nuoM* (1.74-fold) were compared to the wild-type (WT) for their biofilm viability and intracellular RONS accumulation. Relative to PAW-treated WT, *ΔtrxC* and *ΔnuoM* knockout mutants displayed significantly reduced biofilm viability (P ≤ 0.05) confirming their role in PAW-mediated response. Interestingly, *ΔtrxC* biofilms had the highest intracellular ROS accumulation, as revealed by DCFDA staining after PAW treatment. This study gives a detailed insight into how *E. coli* biofilms respond to oxidative stress induced by PAW. It highlights the significance of superoxide in PAW’s bactericidal effects. Overall, our findings shed light on the specific genes and pathways that help *E. coli* biofilms survive and respond to PAW treatment, offering a new understanding of plasma technology and its anti-biofilm mechanisms.

## 1. Introduction

*Escherichia coli* biofilms readily form on stainless-steel^1, 2^, a material used in tools, equipment, storage containers and surfaces across several important industries and sectors: medical, veterinary, water treatment, engineering, agriculture and food processing. Biofilms, consisting of microbial cell aggregates (e.g., bacteria, fungi, viruses) encased in extracellular polymeric substances, differ from planktonic cells in growth rate, gene expression, physiology and metabolism. Biofilm formation enables *E. coli* to survive and persist despite efforts to eradicate them via conventional antimicrobials (e.g., antiseptics, disinfectants, sanitisers and antibiotics) as well as physical removal methods (e.g., scrapping, scrubbing and sonicating). Biofilms are detrimental to materials and serve as a reservoir for infection and disease outbreaks^1, 2^. The global economic cost for biofilms is significant, estimated at approximately $4 B across multiple sectors, with 66% of the economic burden arising from biofilm-associated corrosion^3^. To mitigate this, novel antimicrobial agents and strategies are urgently required that can effectively target biofilms and their cells.

Plasma-activated water (PAW) is a novel technology harnessing plasma, the fourth state of matter. In this state, matter exists in a partially ionised gas, exhibiting a highly volatile environment of excited molecules, UV particles, electromagnetic field and reactive species^4^. Generating plasma directly in water creates several reactive oxygen and nitrogen species (RONS). In PAW generated with air, several short- and long-lived RONS like superoxide, ozone, hydrogen peroxide, hydroxyl radicals, nitrite and nitrate will form^5, 6^. However, when pure oxygen is utilised as an input gas source, reactive oxygen species (ROS) will predominate^5^. The versatility of PAW is attractive and the composition can be further modified dependent on the reactor type, voltage settings, frequency discharge, or by using alternate liquids (e.g., tap water, saline solution and buffers). RONS are widely accepted as the microbicidal component of PAW, with activity against biofilm and planktonic Gram-negative and Gram-positive bacteria, fungi and viruses^5-9^. PAW targets multiple microbial components such as the membrane, proteins, lipids and nucleic acids^4, 6, 10^. Because of the multiple targets and rapid killing, resistance to PAW is relatively rare^11^. We have previously shown that PAW can enhance the anti-biofilm activity of topical chronic wound antiseptics underscoring its clinical application^6^. We have also demonstrated PAW’s ability to outcompete bleach as a fresh produce sanitiser in the context of the food industry^7^. More recently, we have generated PAW with differing gas input sources (air, argon, nitrogen and oxygen), finding PAW generated with oxygen as the most effective at eradicating *E. coli* biofilms that contaminate stainless-steel surfaces^5^. This study found that superoxide was the key ROS implicated in killing *E. coli* biofilm cells^5^.

Despite PAW’s promise as an antimicrobial agent, research has primarily focused on antimicrobial susceptibility assays reliant on colony counts, observing morphological/architectural changes via microscopy (e.g., scanning electron or confocal laser scanning microscopy), intracellular RONS accumulation assays, or observing protein and DNA release. To date, mechanisms underpinning PAW-induced microbial stress responses via proteomics, metabolomics, DNA microarray, RNA-sequencing and transcriptomic techniques are largely underexamined. Some studies have been conducted in the context of planktonic bacteria (*Bacillus cereus, Pseudomonas aeruginosa* and *E. coli*), finding that plasma exposure results in the high expression of genes and pathways to mitigate oxidative stress, such as SOS regulon, DNA repair genes, genes linked to antioxidant production and transporters^12-14^.

In this study, we examined the genetic stress responses of *E. coli* biofilms formed on stainless-steel surfaces after oxygen-PAW treatment. We investigated how superoxide, and its ROS by-products, drive biological processes for biofilm survival during oxidative stress. *E. coli* biofilms were subjected to sub-lethal PAW treatment, their RNA extracted, and gene expression was evaluated. Knockout mutants in the identified genes were then assessed for viability and intracellular RONS accumulation following PAW treatment to confirm their role.

## 2. Materials and Methods

### 2.1 Strain and Culture Conditions

*Escherichia coli* (ATCC 25922) was routinely maintained on Luria-Bertani (LB) agar and cultured in liquid tryptic soy broth (TSB) media at 37°C^5^. The wild-type (WT) *E. coli* K-12 strain (BW25113) and single-gene knockout mutants Δ*trxC*, Δ*nuoM* and Δc*ysP* were utilised from the Keio collection and were cultured with TSB-kanamycin (25 µg/mL)^15^.

### 2.2 Biofilm Formation

Overnight *E. coli* cultures were diluted to ≈ 2×10^7^ CFU/mL, with 1 mL of the diluted culture inoculated into wells of a 24-well plate containing sterile stainless-steel coupons with a diameter of 12.7 mm and thickness of 3.8 mm (BioSurface Technologies, USA)^5^. Biofilms were grown at 30°C with shaking (110 rpm) for 48 h.

### 2.3 Plasma-Activated Water Generation and Treatment

The plasma-activated water (PAW) was generated using conditions and apparatus as previously described^5^. Briefly, using a Leap100 high voltage, microsecond pulsed power source (PlasmaLeap Technologies, Australia), 100 mL autoclave sterilised MilliQ PAW was generated using an input voltage of 150 V, discharge frequency of 1500 Hz, resonance frequency of 60 kHz, and a duty cycle of 100 µs. Oxygen was used as input gas to the plasma generator at a flow rate of 1 standard litre per minute (slm). The biofilms grown on the stainless-steel coupons were placed directly into the Schott bottle and *in situ* PAW-treated (2 min sub-lethal treatment, ≈ 20% biofilm cell death) (Fig. 1). To assess the impact of superoxide on *E. coli* biofilms, 20 mM Tiron (disodium 4,5-dihydroxybenzene-1,3-disulfonate), a superoxide scavenger, was added to the MilliQ prior to PAW generation. Controls of 100 mL autoclave sterilised MilliQ with (Bubble) or without Tiron (Bubble-Tiron) were subjected to 2 min exposure to oxygen flow at 1 slm without plasma discharge.

**Figure 1:**
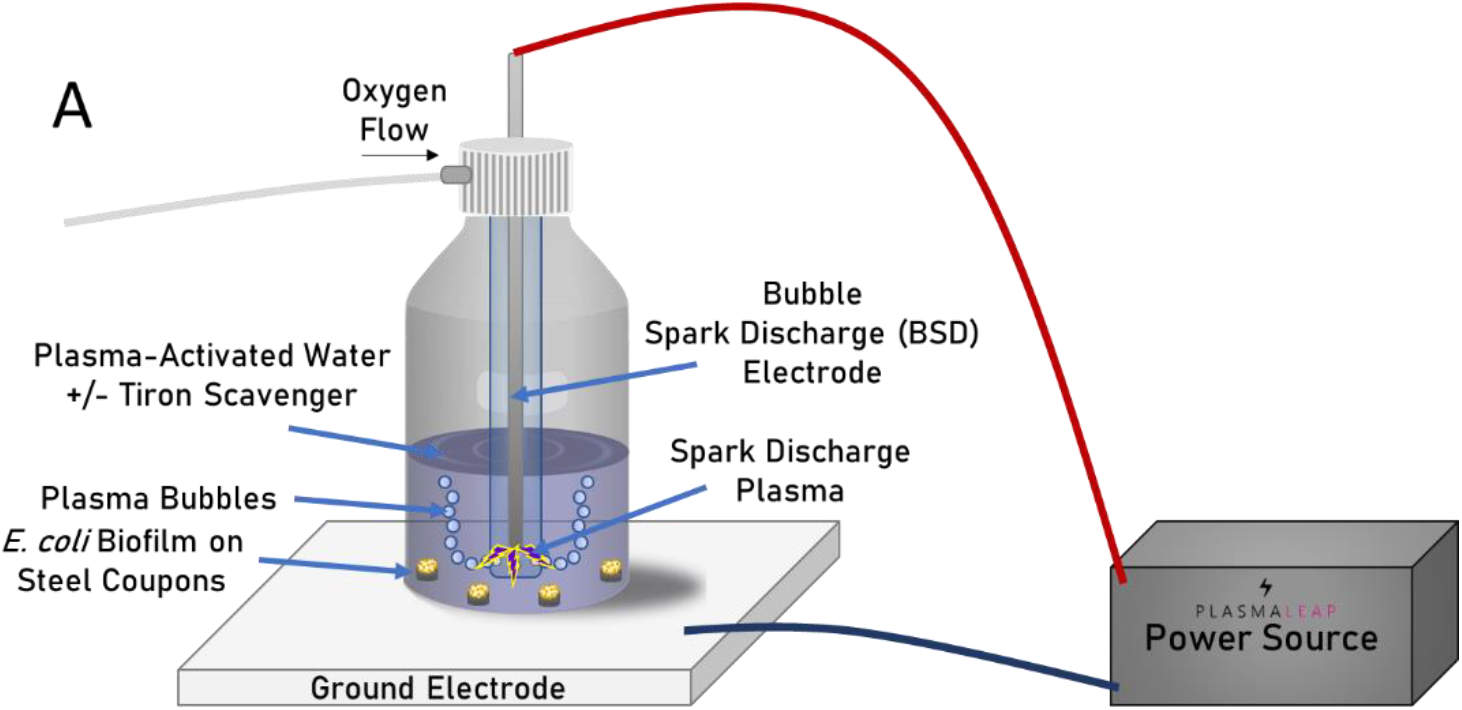
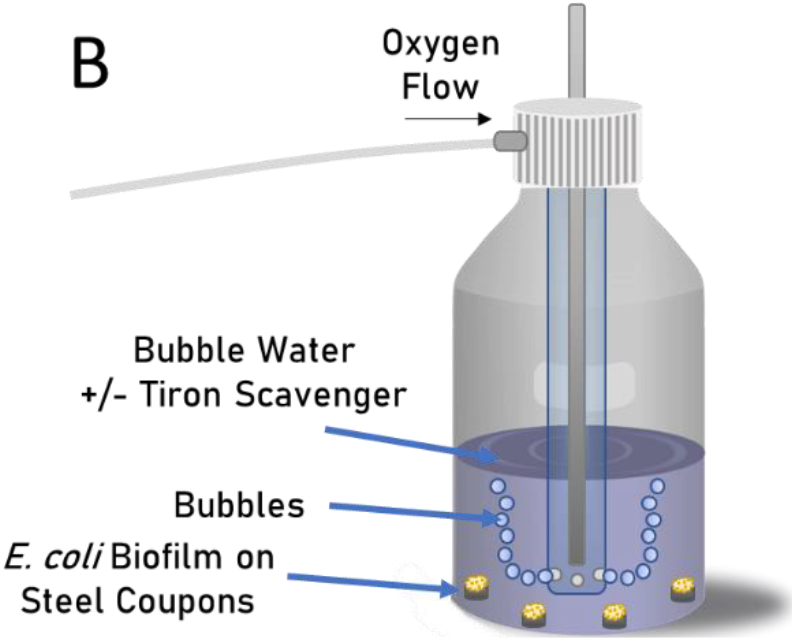
Schematic representation of the reactor utilised to generate PAW and Bubble water used to treat 48 h *E. coli* biofilms grown on stainless-steel coupons. **(A)** A bubble spark discharge (BSD) reactor was used to generate PAW with oxygen as the gas source at 1 slm. PAW was generated with and without 20 mM Tiron scavenger to assess the impact of superoxide anions on biofilm cell stress responses. **(B)** Bubble water controls with and without 20 mM Tiron scavenger was generated by simply passing oxygen through the water at 1 slm without plasma discharge. In both instances, biofilms grown on stainless-steel coupons were treated *in situ* for 2 min. All ATCC 25922 *E. coli* biofilm cells were harvested, and the RNA extracted, sequenced and analysed. Biofilm viability was assessed for Keio single-gene knockout mutants (Δ*trxC*, Δ*nuoM* and Δc*ysP*) and WT harvested from the treated coupons.

### 2.4 PAW Physicochemical Analysis

The physicochemical properties of the PAW, PAW-Tiron, Bubble and Bubble-Tiron treatments including temperature, pH, oxidation-reduction potential (ORP), electrical conductivity, and ozone were measured as described elsewhere^5, 6^. Briefly, a double junction, gel-filled pH probe with built-in temperature sensor was used to measure the pH, ORP was measured using a combination ORP electrode and general-purpose reference electrode, conductivity was measured via a four-ring electrical conductivity probe. These probes and the research-grade benchtop meter were sourced from Hanna Instruments (USA). Dissolved ozone concentrations were determined using a colorimetric assay using the N, N-diethyl-p-phenylenediamine method (accurate at 0.00–2.00 mg/L) with the intensity of the solution at 525 nm measured by a multiparameter benchtop photometer from Hanna Instruments.

### 2.5 Quantification of Intracellular Biofilm Reactive Oxygen and Nitrogen Species

The 2′,7′–dichlorofluorescin diacetate (DCFDA; SigmaAldrich, Australia) and 4,5-diaminofluorescein diacetate (DAF-FM; Sigma-Aldrich, Australia) staining assay was applied to evaluate the accumulation of intracellular ROS and reactive nitrogen species (RNS), respectively^6^. Briefly, *E. coli* biofilms were challenged for 2 min with PAW, PAW-Tiron and controls (Bubble and Bubble-Tiron). The challenged biofilms were then stained with either 20 µM DCFDA or 5 µM DAF-FM for 30 min and measured by CLARIOStar plate reader at Ex/Em of 485-15 nm/535-15 nm or 495-15 nm/515-15 nm, respectively.

### 2.6 RNA Extraction and Sequencing

Four independent biological replicates of 48 h ATCC 25922 *E. coli* biofilms were generated, each comprising four technical replicates that were pooled. Pooled biofilms were treated *in situ* for 2 min with four different treatment solutions PAW, PAW-Tiron, Bubble and Bubble-Tiron. Treatment solutions were gently decanted from the Schott Bottles and the treated biofilm coupons extracted and placed into falcon tubes containing 2 mL sterile 1×PBS. Biofilm cells were dislodged from the coupon surface by 10 sec of vortexing. Lysozyme and proteinase K were utilised to enzymatically lyse and digest the *E. coli* biofilm cells, and RNA extracted as per RNeasy^®^ Mini Kit manufacturer’s protocol (Qiagen). The library preparation and ribosome depletion were carried out using the Illumina Stranded Total RNA prep Ligation with Ribo-Zero Plus kit according to the manufacturer’s protocol (Illumina, San Diego, USA). The libraries were sequenced on the Illumina NextSeq1000 platform (1×100 bp) at the Ramaciotti Center for Genomics (UNSW, Australia).

### 2.7 Transcriptomic Analysis

The quality of RNA-seq reads were checked using FastQC^16^. The sequencing reads were filtered to remove adapter sequences and low-quality bases (≤Q30) using TrimGalore. Quality filtered reads were aligned to the reference genome of *E. coli* ATCC 25922 (RefSeq accession numbers NZ_CP009072) using Subread aligner ^17^. The gene counts per mapped reads were quantified using featureCount^18^. The count matrices were used for differential expression and multi-dimensional scaling (MDS) analysis using edgeR in R version 4^19^. A heatmap of the top 80 differentially expressed transcripts was plotted using mean-centered log2 transformed expression values measured in RPKM (reads per kilobase of transcript per million reads mapped) unit and visualised by pheatmap package in R^20^. The pathway enrichment analysis was performed using ClusterProfiller package in R^21^. The Gene Ontology (GO) enrichment analysis of the upregulated differentially expressed genes (DEGs) was performed in GO web server (http://geneontology.org/) using panther databases (http://www.pantherdb.org) for classification of the genes^22, 23^. Volcano plots and other visualisations were generated with ggplot2 package in R^24^.

### 2.8 Statistical Analysis

All statistical analyses were performed using GraphPad Prism (version 8.4.0, GraphPad Software, USA). A one-way ANOVA was performed with a Tukey’s multiple comparisons post hoc test, and P ≤ 0.05 was considered significant. Fischer’s exact test with FDR (False discovery rate) correction was applied for GO overrepresentation analysis.

## 3. Results and Discussion

PAW contains various RONS crucial for its antimicrobial effectiveness against bacteria such as *E. coli, S. aureus* and *Listeria monocytogenes*^5-7^. PAW, produced with different gas sources, yields diverse RONS^4, 5^. We have previously shown that 10 min oxygen-generated PAW demonstrated superior killing efficacy against *E. coli* biofilms treated *in situ* when compared to PAW generated with other gas input sources (air, nitrogen or argon)^5^. Superoxide, among other ROS (e.g., ozone, hydrogen peroxide, hydroxyl radicals) identified in the PAW, played a pivotal role in its antimicrobial activity^5^ and *E. coli* biofilm viability is improved upon the scavenging of superoxide from PAW with Tiron^5^. Here, biofilms were exposed to PAW treatment for 2 min, which was deemed sub-lethal as this led to ≈ 20% of *E. coli* biofilm cell death compared to the Bubble control (Fig. 2). In contrast, biofilms treated for 2 min with PAW-Tiron (superoxide scavenger), did not significantly impact biofilm viability when compared to Bubble-Tiron and Bubble controls (Fig. 2). A comprehensive assessment of the physicochemical properties of the PAW that was generated for 2 min was performed (Supplementary Table. 1). Similar to previously analysed oxygen-PAW that was generated for 10 min, our analysis confirmed the presence of ozone (a precursor of superoxide), hydrogen peroxide, hydroxyl radicals, and other ROS by-products. As supported by our high oxidation-reduction potential values (Supplementary Table. 1), ROS concentrations were notably higher in PAW (514.33 ± 12.50 mV) than PAW-Tiron (279.63 ± 3.13 mV) and controls (Bubble, 190 ± 2.00 mV and Bubble-Tiron, 281.43 ± 1.25 mV). These results are comparable with our previous study^5^.

**Figure 2:**
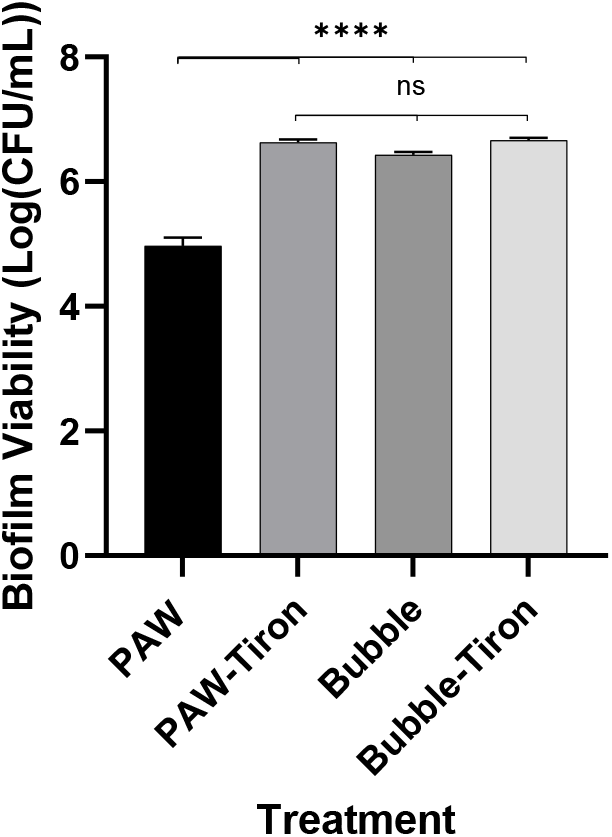
Two min *in situ* PAW treatment reduces *E. coli* biofilm viability by ≈20% when compared to PAW-Tiron and controls (Bubble and Bubble-Tiron) Data represents mean ± standard error of the mean, **** (P ≤ 0.0001); non-significant (P > 0.05); n = 3 biological replicates, with 2 technical replicates each.

PAW is abundant in ROS that can severely damage microbial structures such as membranes, lipids, enzymes, proteins and nucleic acids^6, 25^. Previous research suggests that regulatory systems (e.g., SoxRS and OxyR) and DNA repair processes play pivotal roles in mitigating these effects^26-28^. To ensure survival, bacteria can up-or down-regulate genes to enhance viability. Understanding the mechanisms by which *E. coli* respond to PAW-induced oxidative stress is crucial^26, 27^. To explore the oxidative stress responses of *E. coli* biofilms to sub-lethal PAW exposure, we conducted RNA-seq analysis followed by transcriptomic analysis to assess gene expression changes.

Analysis revealed a marked difference in gene expression in *E. coli* biofilms formed on stainless-steel surfaces when exposed *in situ* to sub-lethal 2 min PAW treatment as depicted by MDS plot (Fig. 3A) and clustering heatmap of top differentially expressed genes (Supplementary Fig. 1). A strong correlation was also observed among the RNA-seq data for biological replicates of PAW, PAW-Tiron, Bubble and Bubble-Tiron treatment groups. Around 40% of the variation in gene expression was observed for *E. coli* biofilms treated with PAW compared to PAW-Tiron and controls (Bubble and Bubble-Tiron). PAW treatment resulted in 478 upregulated genes (> 1.5 log2FC) and 186 downregulated genes (< −1.5 log2FC) compared to Bubble control treated biofilm cells (Fig. 3B). The removal of superoxide from PAW (i.e., PAW-Tiron treatment) resulted in relatively fewer upregulated genes compared to controls (99 genes Bubble control, Fig. 3D, and 91 genes Bubble-Tiron control, Fig. 3E). The implications of superoxide on *E. coli* gene expression were further supported by comparing PAW and PAW-Tiron (Fig. 3F) treatments, resulting in 453 upregulated genes. Lastly, there were no significant differences in gene expression between Bubble and Bubble-Tiron treatments, indicating that the Tiron scavenger does not impact *E. coli* biofilm gene expression (Fig. 3A and Supplementary Fig. 1 and 2).

**Figure 3:**
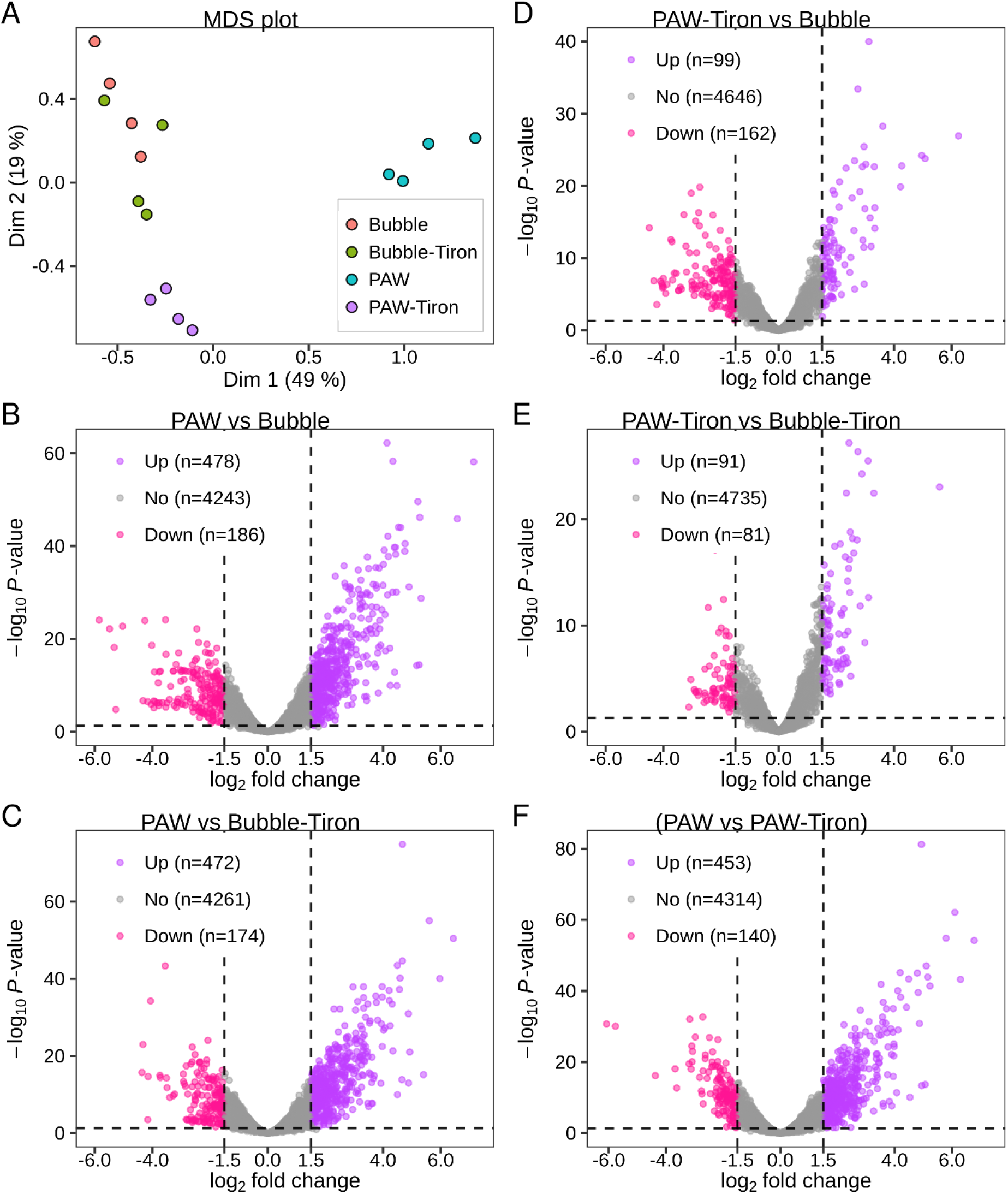
Differential gene expression analysis of *E. coli* biofilms during 2 min *in situ* treatment with PAW, PAW-Tiron, Bubble and Bubble-Tiron. **(A)** MDS plot depicting distances between transcript expression profiles of *E. coli* biofilms treated with PAW (Cyan) compared to PAW-Tiron (Violet) and controls Bubble (Red) and Bubble-Tiron (Green). **(B-F)** The volcano plot represents the differentially expressed transcripts between PAW vs Bubble **(B)**, PAW vs Bubble-Tiron **(C)**, PAW-Tiron vs Bubble **(D)**, PAW-Tiron vs Bubble-Tiron **(E)** and PAW vs PAW-Tiron **(F)** groups as depicted in the title. The log-fold change (base 2) is plotted on the x-axis and the negative log of P-value (base 10) is plotted on the y-axis. Data shows individual Log FC changes in expression between pooled *E. coli* biofilms (4 biological replicates, each comprising 4 pooled technical replicates). Up- and down-regulated genes are represented by violet and pink circles, respectively, while non-significant genes are represented by grey circles.

Given that the *E. coli* biofilm viability data (Fig. 2), differential gene expression analysis (Fig. 3A), heatmap (Supplementary Fig. 1) and linear regression analysis (Supplementary Fig. 2) revealed similarities between Bubble and Bubble-Tiron, these were combined and used for comparison to PAW treatment in pathway enrichment and biological process enrichment analysis (Fig. 4). In *E. coli* biofilms treated with PAW vs control (combined Bubble treatment) (Fig. 4A), sulfur metabolism rich factor (0.4) was the highest compared to any other pathway. Microorganisms require sulfur to function and have taken advantage of its unique characteristics in their metabolic processes: metal binding, nucleophilicity, disulfide bond strength and redox capacity^29^. Sulfur is present in several essential enzyme co-factors and in proteins via amino acids methionine and cysteine. If environmental sulfur is not available, cystine is rapidly reduced to cysteine at the expense of antioxidants (e.g., thioredoxin and glutathione), mediating *E. coli* metabolic stress. This process requires a delicate balance between metabolic need and risk as it can result in cysteine accumulating within *E. coli*^29^. Excess cysteine sensitises the cell to hydrogen peroxide, arising from superoxide. Hydrogen peroxide is disruptive to iron-containing enzymes via Fenton chemistry, and the resultant hydroxyl radicals damage *E. coli* DNA^30, 31^. To counter this, *E. coli* can export cysteine via the L-alanine exporter, AlaE^32^. Interestingly, the *alaE* gene was significantly upregulated in our PAW treated *E. coli* biofilm cells. Moreover, superoxide is also detrimental to iron-sulfur clusters which are valuable to cells: acting as catalysts, structural elements and redox sensors^28, 33^. In this scenario, *E. coli* prioritises iron-sulfur cluster repair^28^. In our PAW treated biofilm cells, *ytfE*, a gene that encodes the iron-sulfur cluster repair protein YtfE, was significantly upregulated. Conversely, when superoxide was removed via the Tiron scavenger (PAW-Tiron treatment), *ytfE* was not significantly upregulated, confirming that superoxide, and its resulting ROS, are key proponents of PAW-induced oxidative stress.

**Figure 4:**
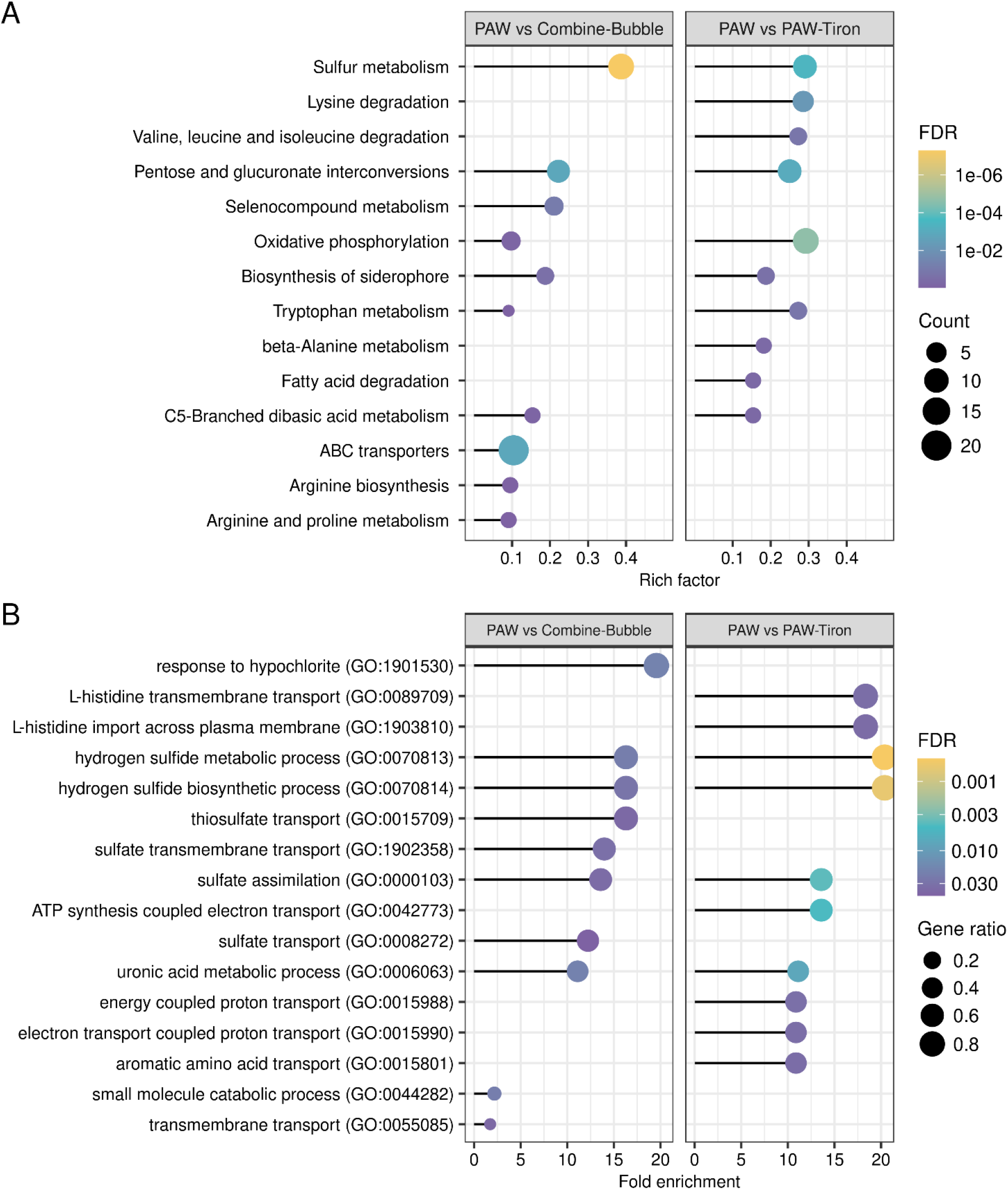
Pathway and GO enrichment analysis of the upregulated differentially expressed genes. **(A)** KEGG pathway analysis of the upregulated DEGs in PAW treatment compared to the control conditions. The size of each dot is proportional to the number of upregulated DEGs for the given pathway in the reference list. Only pathways with rich factor more than 1 are shown for simplicity. **(B)** The top GO terms annotated in the biological process category for the upregulated genes are selected based on their fold-enrichment values. GO terms with fold-enrichment value more than 1 are shown only for simplicity. Fold-enrichment values were calculated from the number of genes observed in the upregulated DEGs list divided by the expected number in the reference list for a particular GO term. The size of each dot is proportional to the number of upregulated DEGs for the given GO term in the reference list. Colour bar representing false discovery rate.

Similarly, genes involved in hydrogen sulfide metabolic/biosynthetic processes and sulfate/thiosulfate transport systems were notably upregulated (fold enrichment values ranging 12.5-17) (Fig. 4B). Over a dozen ATP-binding cassette (ABC) transporter genes were significantly upregulated in PAW vs Bubble treatment (rich factor 0.1, Fig. 4A). ABC transporters require ATP or ADP to actively transport molecules (e.g., ions, amino acids, sugars, lipids, antibiotics) across the lipid membrane to ensure secretion of toxins and uptake of nutrients^34^. In Gram-negative bacteria, the ABC transporter complex comprises of a periplasmic substrate binding protein, two transmembrane domains and two nucleotide-binding domains^35^. In *E. coli*, sulfate assimilation is initiated by sulfate-binding and thiosulfate-binding proteins (encoded by *sbp* and *cysP*, respectively) with overlapping functions in the sulfate/thiosulfate ABC transporter^36^. *sbp* and *cysP* were significantly upregulated upon PAW treatment (1.69- and 1.58-fold, respectively), and are linked to mitigating oxidative stress in *E. coli*^35^. Moreover, once thiosulfate enters the cell, it is converted to S-sulfocysteine, which is then converted to cysteine via antioxidants, thioredoxin (encoded by *trxC*) and glutaredoxin (encoded by *grxA*)^37^. Cysteine is also a major determinant of synthesising another crucial antioxidant, glutathione (encoded by *gorA*). Thioredoxin, glutaredoxin and glutathione can protect *E. coli* against oxidative damage by reducing sulfenic acids and disulfide bonds^28^. As demonstrated by our results, *trxC* and *grxA*, positively regulated by OxyR under hydrogen peroxide and superoxide stress conditions^38^, was significantly upregulated during PAW treatment (4.23- and 2.21-fold, respectively). Conversely, the PAW-Tiron treatment (PAW without superoxide) did not significantly upregulate genes involved in thiosulfate and sulfate transport, and the sulfur metabolism pathway rich factor (0.3) was slightly reduced (Fig 4.A) demonstrating the importance of superoxide in the process.

Other ABC transporter genes were significantly upregulated including *mdlA* (encoding a putative multidrug efflux pump)^39^ and *ribsD/A/C* (encoding ribose transport proteins of the inner membrane)^40^. ABC transporters also process amino acids, it was unsurprising that several amino acid metabolic/biosynthesis (e.g., arginine and proline) pathways (Fig. 4A) were significantly upregulated (rich factor 0.1) in PAW vs combined Bubble Treatment. However, after scavenging superoxide radicals (PAW vs PAW-Tiron), pathways associated with aromatic amino acids (e.g., histidine, phenylalanine, tryptophan and tyrosine) predominated (Fig 4.B). Amino acids found in proteins are the primary sites for oxidative damage^33^. Damage to methionine and cysteine are considered reversible, whilst damage to amino acids like arginine, proline, histidine and tyrosine is irreversible^33^. Studies have suggested that bacteria have adapted the ability to assemble amino acids to counter ROS effects. For example, prioritising methionine/cysteine on the surface of cytosolic proteins can act as “sponges” to absorb oxidative damage^41, 42^. This may explain the differential upregulation in amino acid pathways, particularly when superoxide (and its by-product ROS) is removed.

Superoxide in PAW will readily lead to the formation of hydrogen peroxide that will convert to hypochlorite within *E. coli* cells. This explains the significantly enriched hypochlorite response systems (fold enrichment value of 20) seen only for PAW vs Combined Bubble treated biofilms (Fig. 4B). Lastly, it is unsurprising that genes involved in the oxidative phosphorylation pathway necessary for defence against ROS, were also significantly upregulated (Fig. 4A). Of note, multiple genes encoding the 13 subunit NADH:ubiquinone oxidoreductase complex were upregulated. NADH:ubiquinone oxidoreductase (complex I) is the first of six major cytoplasmic membrane enzyme complexes catalysing oxidative phosphorylation. In humans, complex I deficiencies are linked to several diseases (e.g., Parkinsons) and the aging process because of superoxide and hydrogen peroxide formation within mitochondria. The marked similarities (and differences) between human and *E. coli* complex I have led to insights on how complex I reduces oxygen, generating superoxide and hydrogen peroxide^43^. Complex I is a 13-subunit complex comprising of the membrane arm (subunits NuoA, H, J, K, L, M and N) and peripheral arm (subunits NuoB, CD, E, F, G and I)^44^. Complex I oxidises NADH, reduces ubiquinone, and maintains the proton motive force of the membrane^43, 45^. NuoM and NuoL comprise half the membrane arm of complex I and contain ubiquinone binding sites. Ubiquinone can function as an antioxidant in *E. coli*^46^. Our results found that the *nuoM* gene was the most prominent and was significantly upregulated upon PAW treatment (1.74-fold).

*trxC, cysP* and *nuoM* were selected based on their potential role in ensuring *E. coli* biofilm survival under oxidative stress upon PAW treatment. *trxC, cysP* and *nuoM* genes encode thioredoxin, thiosulfate/sulfate ABC transporter substrate-binding protein CysP, and NADH-quinone oxidoreductase subunit M, respectively. To assess the impact of *trxC, nuoM* and *cysP* genes, *E. coli* biofilms were cultivated using mutants from the Keio collection. These strains demonstrated similar viability to the reference strain ATCC 25922 following treatments with PAW, PAW-Tiron and controls (Supplementary Fig. 3). Both WT and single-gene knockout mutants (*ΔtrxC, ΔcysP* and *ΔnuoM*) were evaluated for biofilm viability after 2 min PAW treatment, compared with PAW-Tiron and control groups (Bubble and Bubble-Tiron) (Fig. 5A). Moreover, ozone and superoxide (and its by-products hydrogen peroxide and hydroxyl radicals) can cause membrane disruption which leads to ROS penetrating the cells. Once within the cells, the ROS can trigger a cascade of oxidation-reduction reactions via Fenton reaction, iron-sulfur clusters, and flavoproteins. Additional potent ROS (e.g., short-lived hydroxyl radical species) are produced and accumulate intracellularly, causing significant damage^25, 47^. Hence, DCFDA and DAF-FM staining was also utilised to detect intracellular ROS and RNS biofilm accumulation, respectively (Fig. 5B and C).

**Figure 5:**
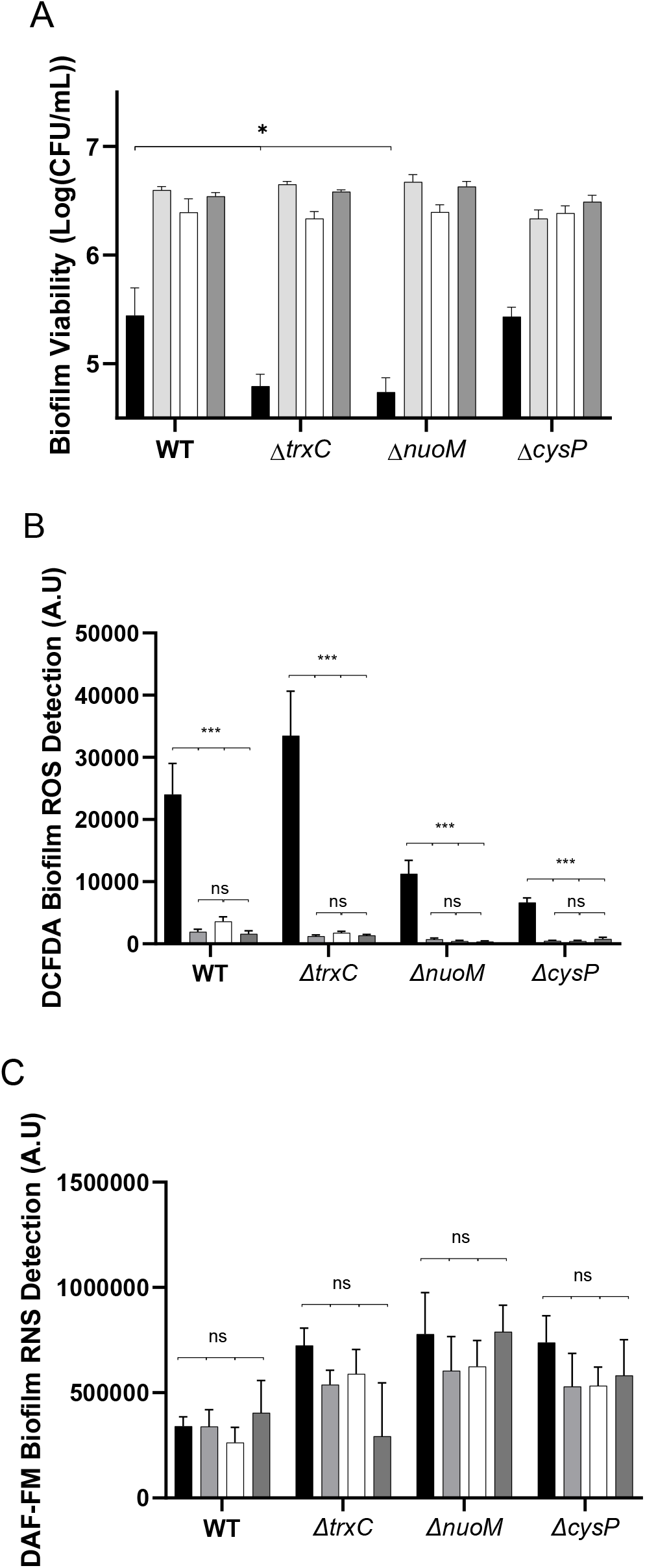
Two min PAW treatment significantly impacts *E. coli* biofilm viability and causes intracellular ROS accumulation. **(A)** Effect on biofilm viability of PAW (black), PAW-Tiron (light grey), Bubble (white), and Bubble-Tiron (dark grey) treatments on *E. coli* biofilm WT and gene knock out mutants Δ*trxC*, Δ*nuoM* and Δ*cysP* was investigated. **(B)** DCFDA staining was used to detect and measure intracellular biofilm ROS accumulation whilst RNS were stained with **(C)** DAF-FM. Data represents mean ± SEM, * (P ≤ 0.05); *** (P ≤ 0.001); ns (P > 0.05); n = 3 biological replicates, with 2 technical replicates each.

Relative to PAW-treated WT biofilms, both Δ*trxC* and Δ*nuoM* mutants exhibited significantly decreased biofilm viability showing that these genes are important in the biofilm cells defence against the ROS found in PAW. Similarly, intracellular ROS accumulation was significant in biofilms of both mutants after PAW treatment. Interestingly, intracellular ROS levels in the PAW treated *ΔtrxC* biofilms surpassed WT levels. These results underscore the importance of thioredoxin’s antioxidant activity. Moreover, it suggests that the inability to convert S-sulfocysteine to cysteine via thioredoxin is detrimental to *E. coli* under PAW’s highly oxidative stress conditions. Additionally, the NuoM subunit which facilitates the binding of ubiquinone, shown to have some antioxidant properties^46^, may offer some protection against oxidative stress induced by PAW-associated ROS. One study showed decreased NADH:Ubiquinone activity when both NuoM and NuoL subunits were removed from complex I^48^. Further study could harness a *nuoM nuoL* double-mutant to assess *E. coli* survival under the harsh ROS conditions of PAW. Nonetheless, these results indicate that both *trxC* and *nuoM* genes are important in the cells defence against PAW-induced oxidative stress.

Although *ΔcysP* biofilms did not show significantly decreased biofilm viability compared to PAW-treated WT, DCFDA staining revealed significant intracellular ROS accumulation after 2 min PAW treatment (Fig. 5B). As *cysP* and *sbp* have partially overlapping activities in transporting thiosulfate and sulfate, single mutation of just one gene may not significantly inhibit this process^36^. However, double-mutants are unable to utilise sulfate or thiosulfate^36^. Further assessment of a double *cysP* and *sbp* mutant may demonstrate significantly reduced *E. coli* biofilm viability and exacerbate intracellular ROS accumulation.

Lastly, biofilm viability for WT and knockout mutants did not significantly differ between PAW-Tiron and controls (Bubble and Bubble-Tiron), suggesting that the removal of superoxide via Tiron scavenger in PAW significantly lowers PAW’s antibacterial activity and promotes biofilm survivability. Intracellular RNS was detectable within *E. coli* biofilms treated with all four treatments (PAW, PAW-Tiron, Bubble and Bubble-Tiron), albeit in slightly elevated levels for mutant strains (Fig. 5C). However, there were no significant differences in intracellular RNS accumulation between treatment types.

## 4. Conclusions

This study presents, for the first time, new insights into PAW’s mechanisms of action. Providing novel knowledge of the genetic stress responses *E. coli* biofilms evoke under oxidative stress conditions. Within just 2 min of PAW treatment, biofilm viability is significantly decreased, and a complex cellular response is prompted by upregulating 478 genes. These results underscore PAW’s utility as an anti-biofilm agent. Further work is warranted to better understand long-term effects over multiple generations of *E. coli* biofilm bacteria. Assessing a broader range of single- and double-gene knockout mutants of other genes we found significantly upregulated (e.g., *alaE, ytfE, sbp*) would greatly improve our understanding of PAW. These findings prompt investigation into further customising PAW, exploring different treatment times, changing oxygen flow rates, and altering voltages to assess and compare consequent genetic responses.

## 5. Acknowledgements

This work was supported by the Australian Research Council Discovery Scheme (DP210101358). We thank A/Prof Amy Cain (Macquarie University) for kindly providing the Keio collection strains utilised in this study. We thank Prof Dee Carter (University of Sydney), Prof Diane McDougald (University of Technology Sydney), Dr Nick Coleman (Macquarie University) and Dr Behdad Soltani (University of Sydney) for their invaluable support of this project. The authors acknowledge the Ramaciotti Centre for Genomics for their assistance.

## 6. Author contributions

Conceptualisation, H.K.N.V. and A.M-P.; methodology, H.K.N.V.; formal analysis, H.K.N.V., M.M.H., and B.X.; experimental investigation, H.K.N.V., B.X., and D.A.; data curation, H.K.N.V., M.M.H., and B.X.; visualisation, H.K.N.V.; writing—original draft preparation, H.K.N.V.; writing— review and editing H.K.N.V., M.M.H., B.X., D.A., P.J.C., S.A.R., and A.M-P.; supervision, A.M-P. and H.K.N.V.; resources, A.M-P., S.A.R., and P.J.C.; funding acquisition, A.M-P. All authors have read and agreed to the published version of the manuscript.

## Supplementary

**Table 1:**
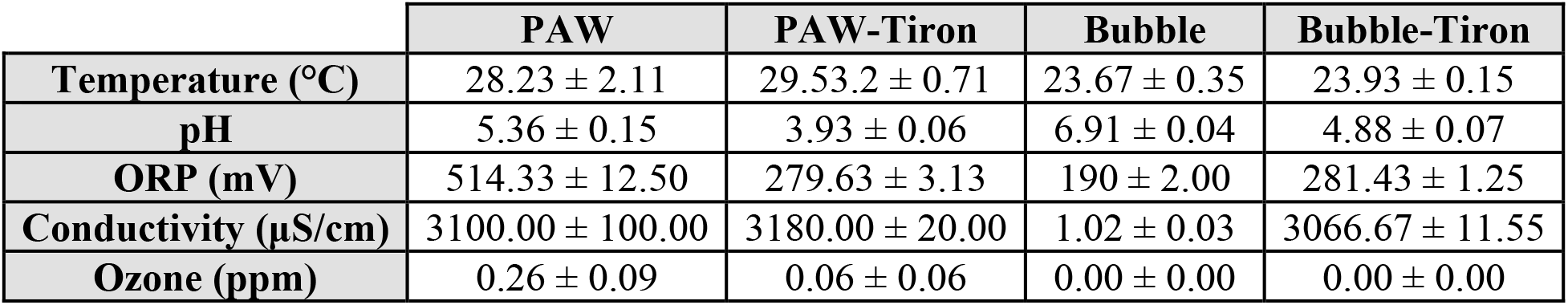
The physicochemical properties of PAW, PAW-Tiron and controls (Bubble and Bubble-Tiron) generated for 2 min. Data represents mean ± standard deviation, n = 3 replicates.

|  | PAW | PAW-Tiron | Bubble | Bubble-Tiron |
| --- | --- | --- | --- | --- |
| Temperature (°C) | 28.23 ± 2.11 | 29.53.2 ± 0.71 | 23.67 ± 0.35 | 23.93 ± 0.15 |
| pH | 5.36 ± 0.15 | 3.93 ± 0.06 | 6.91 ± 0.04 | 4.88 ± 0.07 |
| ORP (mV) | 514.33 ± 12.50 | 279.63 ± 3.13 | 190 ± 2.00 | 281.43 ± 1.25 |
| Conductivity (µS/cm) | 3100.00 ± 100.00 | 3180.00 ± 20.00 | 1.02 ± 0.03 | 3066.67 ± 11.55 |
| Ozone (ppm) | 0.26 ± 0.09 | 0.06 ± 0.06 | 0.00 ± 0.00 | 0.00 ± 0.00 |

**Figure 1:**
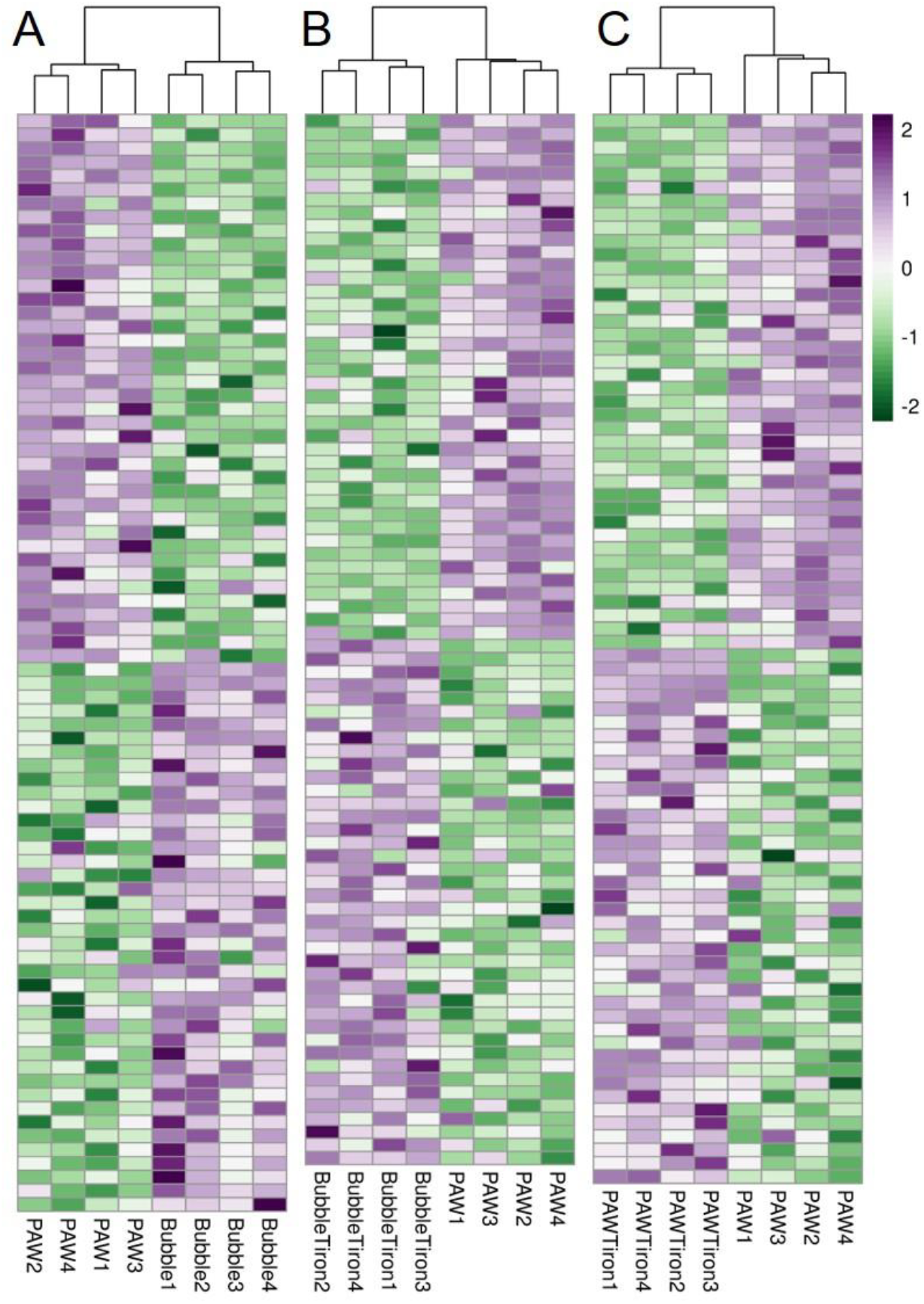
Heatmap of RNA-seq data showing top 80 differentially expressed transcripts (40 from each up and down-regulated groups) in *E. coli* biofilms treated with PAW vs Bubble (A), PAW vs Bubble-Tiron (B) and PAW vs PAW-Tiron (C) Data represent the mean-centred log2 transformed expression values measured in RPKM units (reads per kilobase of transcript per million reads mapped). The dendrogram represents clustering of biological replicates based on their expression profile.

**Figure 2:**
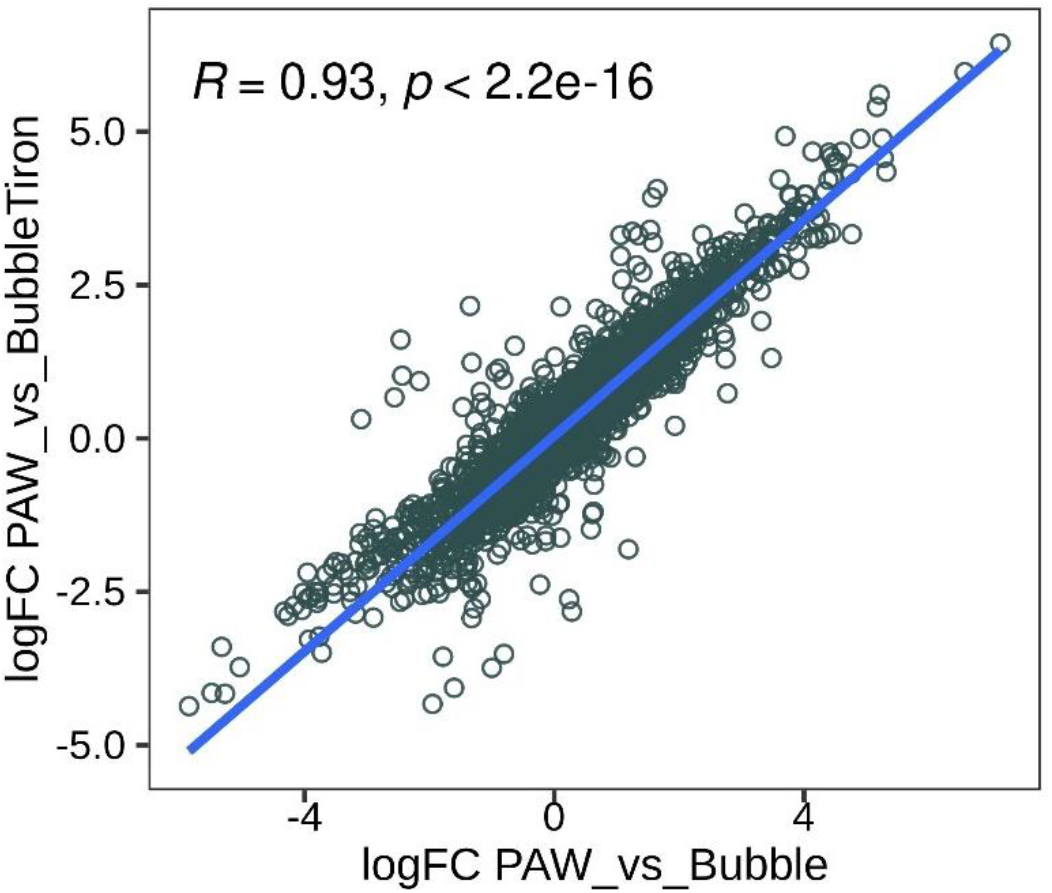
Linear regression analysis was applied after differential expression for *E. coli* biofilms treated for 2 mins with PAW compared to Bubble-Tiron and PAW compared to Bubble. This data confirms statistically significant similarity between Bubble controls.

**Figure 3:**
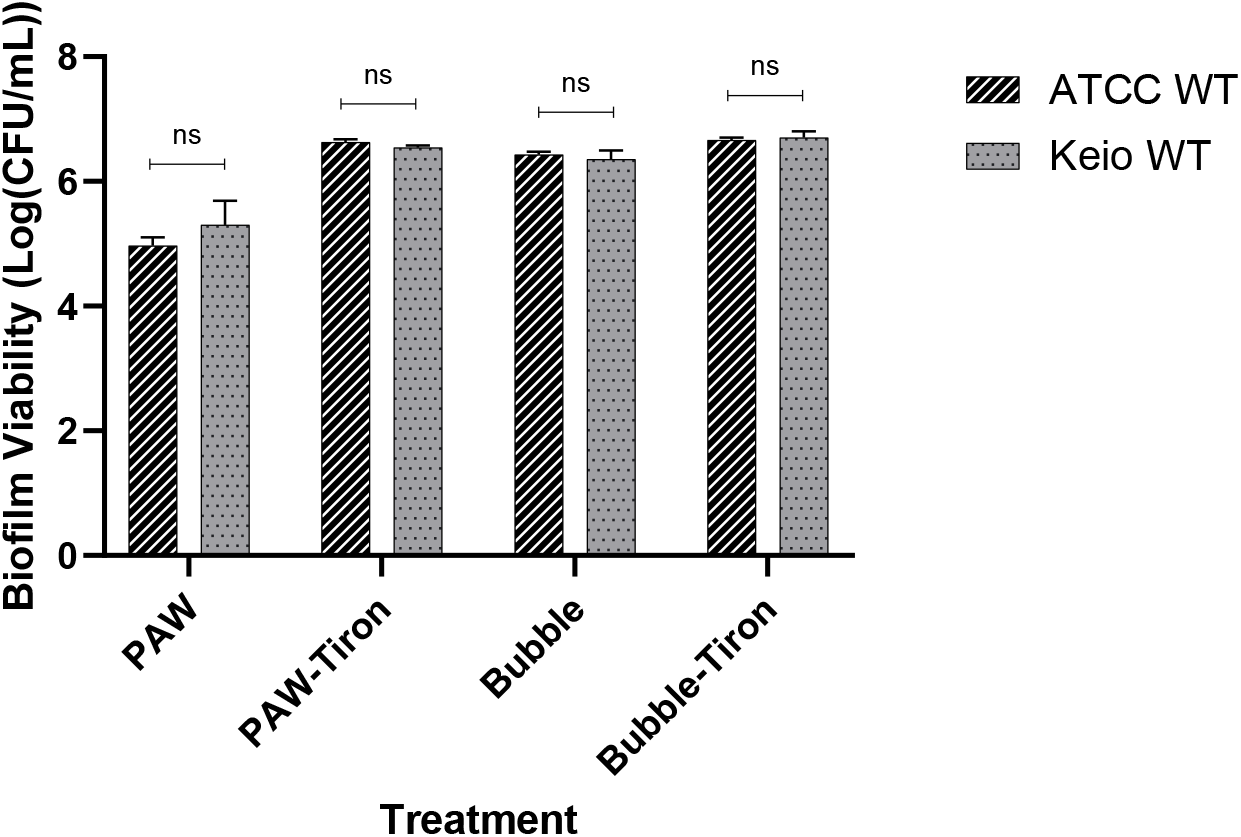
Biofilm viability of *E. coli* WT ATCC and Keio Collection strains demonstrating phenotypic similarity after 2 min PAW, PAW-Tiron, Bubble and Bubble-Tiron treatments. This data validates usage of WT and single-gene knockout mutants from the Keio collection for viability and intracellular RONS assessment in Figure 5. Data represents mean ± SEM, ns (P > 0.05); n = 3 biological replicates, with 3 technical replicates each.

